# Population coding of grasp and laterality-related information in the macaque fronto-parietal network

**DOI:** 10.1101/179184

**Authors:** Jonathan A Michaels, Hansjörg Scherberger

## Abstract

Preparing and executing grasping movements demands the coordination of sensory information across multiple scales. The position of an object, required hand shape, and which of our hands to extend must all be coordinated in parallel. The network formed by the macaque anterior intraparietal area (AIP) and hand area (F5) of the ventral premotor cortex is essential in the generation of grasping movements. Yet, the role of this circuit in hand selection is unclear. We recorded from 1342 single- and multi-units in AIP and F5 of two macaque monkeys (Macaca mulatta) during a delayed grasping task in which monkeys were instructed by a visual cue to perform power or precision grips on a handle presented in five different orientations with either the left or right hand, as instructed by an auditory tone. In AIP, intended hand use was only weakly represented during preparation, while hand use was robustly present in F5 during preparation. Interestingly, visual-centric handle orientation information dominated AIP, while F5 contained an additional body-centric frame during preparation and movement. Together, our results implicate F5 as a site of visuo-motor transformation and advocate a strong transition between hand-invariant and hand-specific representations in this parieto-frontal circuit.

## Introduction

Our everyday reaching and grasping movements demand the coordination of information across multiple scales. While grasping a cup requires determination of the physical position and orientation of the cup, one must also resolve the appropriate shaping of the hand, which hand to use, and the muscle forces required. Given this, and given the flexibility with which we switch between hands, it is expected that both hand independent and muscle specific representations should be found at various levels of abstraction throughout cortex.

Indeed, a number of studies have probed how neural circuits represent laterality of reaching movements in macaque monkeys. Integration of arm specific and arm independent information has been found in the posterior parietal cortex^1-3^, premotor cortex^4-9^, and primary motor cortex (M1)^10-12^, although the outputs from M1 have been identified as mostly contra-lateral^13,14^.

Yet, little is known about the laterality of grasping movements. It has been shown that when all inter-hemispheric connections of macaques has been severed, the ipsi-lateral hemisphere can generate reaching movements towards food, but cannot properly pre-shape the fingers of the hand^15^, suggesting that grasping is a highly lateralized process. The hand grasping circuit^16^ consisting of the hand area (F5) of the ventral premotor cortex and the anterior intraparietal area (AIP) is an essential anatomical and functional circuit in grasp preparation and execution. Neural activity in these areas is strongly modulated by visual object properties^17,18^, extrinsic goals^19^, performed grip types^20,21^, and preparatory activity in these areas can be used to decode the visual properties of objects and complex hand shapes required to grasp a diverse range of objects^22-24^, as well as predict reaction times^25^. Although laterality has been studied in ventral premotor cortex, these studies either employed no delay period^26^, simple movements^10^, or required only reaching movements^6,7,27^. Additionally, to our knowledge, no electrophysiological studies of laterality have been undertaken in AIP.

In the current study, laterality of grasping movements were investigated using a delayed grasping task^28^ while neural activity was recorded in AIP and F5. Two monkeys visually fixated a central fixation point throughout the trial. During a cue phase, monkeys received a visual cue indicating which of two grip types to perform in one of five possible grasping handle orientations as well as an auditory tone indicating the hand to use on that trial. Following a memory period, a go cue instructed monkeys to grasp the handle in the dark.

We found that activity in AIP and F5 during the movement robustly reflected which hand was used, but preparatory activity representing the intended hand was mostly found in F5, suggesting that AIP represents task information independent of hand during preparation. Furthermore, the amount of grip tuning and preferred grip type of each unit did not depend on hand used, indicating a shared framework for grasp planning. However, although orientation tuning was abundant in AIP, typically lasting for the entire trial, orientation tuning was present in F5 primarily for contra-lateral movements, revealing a functional differentiation between hemispheres. Crucially, while visual-centric coordinate frames were present in both areas, a body-centric coordinate frame representing handle orientation in a mirror-symmetric fashion was present in F5 starting towards the end of the cue period and lasting throughout the movement, implicating F5 as a site for visuo-motor transformation during movement preparation.

## Results

### Behavior

To investigate the laterality of grasp movement coding in premotor and parietal cortex, two monkeys performed a delayed grasping task in which the hand the monkey had to use, as well as the appropriate grip type and hand orientation, were cued on each trial (Fig. 1a,b). Concurrently with behavior, single- and multi-unit activity was recorded from premotor area F5 and parietal area AIP simultaneously (Fig. 1c,d). Both monkeys successfully performed the task. After initiating trials to the point of obtaining specific trial information, monkeys S and P successfully completed 85% and 84% of trials, respectively. In detail, monkeys S and P correctly selected the correct hand on 89% and 93% of trials, respectively, while grip type selection was correct 99% and 98% of the time. In addition to keeping their hands on the hand rests, all motion of the hand was tightly controlled with a separate tracking program using infrared camera data (Methods). Trials where miniscule hand movement was detected were aborted without reward. Trials were completed successfully without premature movement 99% and 94% of the time, for monkeys S and P, respectively. Median reaction time, i.e. the time between the go cue and the hand leaving the handrest, were 230 and 265 ms for monkeys S and P, respectively, while median movement time, i.e. the time between the hand leaving the handrest and executing the appropriate grip on the handle were 305 and 325 ms. In detail, movement times were 325 and 320 ms for the left and right hands, respectively, for monkey P, and 305 ms for both hands in monkey S.

**Figure 1.**
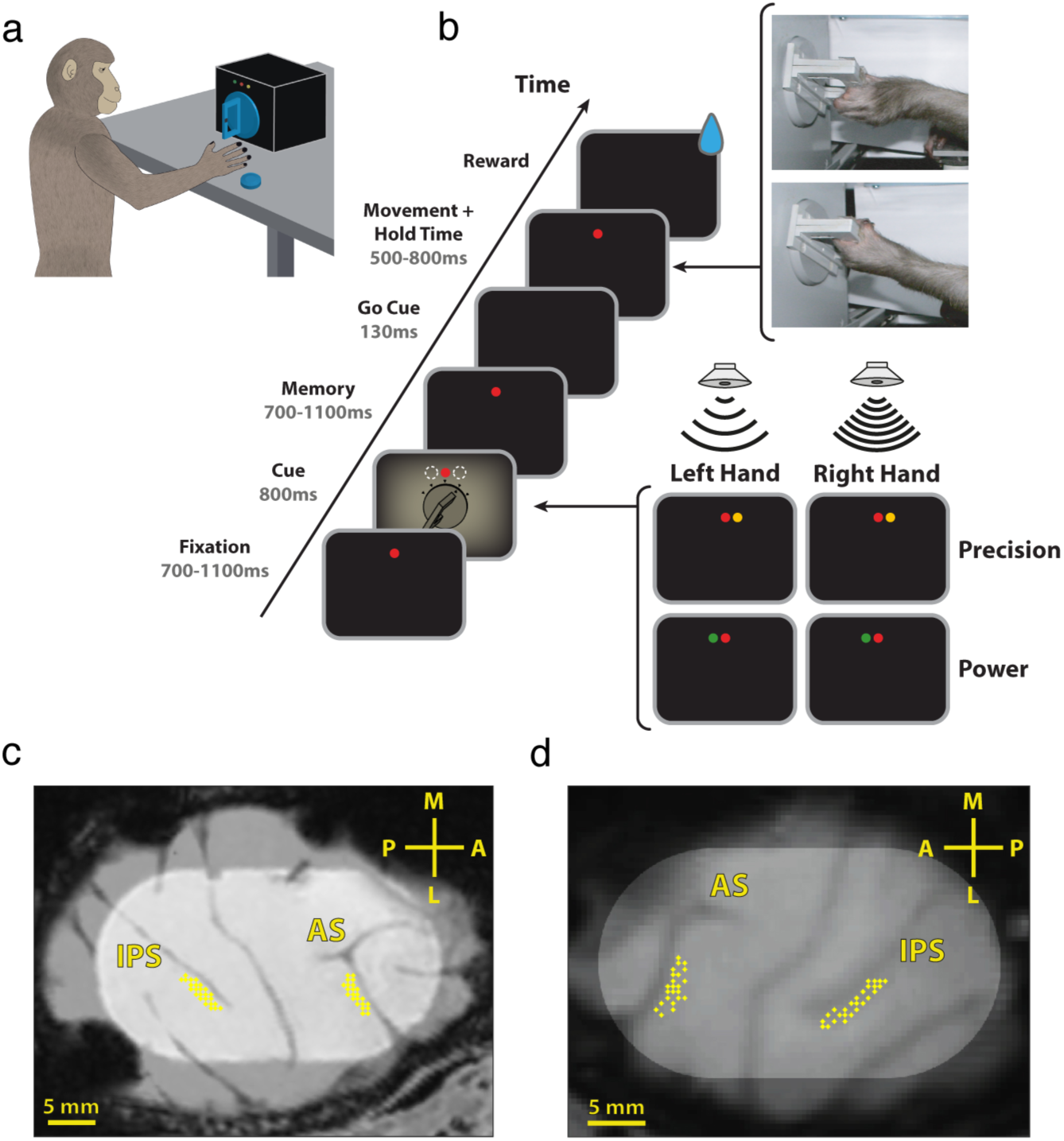
Task design and recordings. (a) Illustration of a monkey in the experimental setup. The cues were presented via LEDs above the handle. (b) Delayed grasping task with two grip types (top: power grip, bottom: precision grip), five orientations of the grasping handle, and grasped with either the left or right hand. Grips and orientation were cued using LEDs and handle illumination, while hand was cued by two auditory tones. Trials were presented in pseudorandom order in darkness. c-d, Recording locations for monkey P (c) and S (d) overlaid on a structural MRI. The illuminated oval marks the outline of the recording chamber. Recordings were made in F5 on the bank of the arcuate sulcus (AS) and in AIP toward the lateral end of the intraparietal sulcus (IPS). The cross shows medial (M), lateral (L), anterior (A), and posterior (P) directions. Note that monkey S was implanted in the left hemisphere and monkey P in the right hemisphere.

### Neural recordings

The analyzed data sets included a collection of 178 individual recording sessions, 91 from monkey S and 87 from monkey P. In monkey S, 861 single- and multi-units were successfully recorded (single: 459, multi: 402), of which 581 were task-related (AIP: 189, F5: 392) and used in further analysis (Methods). In monkey P, 481 units were recorded (single: 263, multi: 218), of which 390 were task-related (AIP: 207, F5: 183). Units were classified as task-related if they were tuned for any of the three task factors (hand, grip, or orientation) at any point during the course of the trial as determined by a cluster-based permutation test (CBPT; Methods), which finds contiguous segments of time tuned for one of the three task factors, while keeping the overall false-positive rate below 5% over all three factors and time points. Qualitatively similar results were obtained using an ANOVA with a sliding window with multiple comparison corrections. Only units found to be task-related were used in further analysis.

To get an overview of what kind of task-related responses were present, we averaged over all trials of each condition to produce average firing rate curves and combined them with the significance testing described above. Figure 2 shows a number of example single-units recorded from both areas and monkeys. One of the most common responses in AIP was tuning for a specific orientation of the handle that was sustained from cue onset to the end of movement, even though the handle was only illuminated during the cue (Fig. 2 - Top Left). Another common response in AIP was units that did not respond to the cue at all, but showed strong grip and hand tuning specifically during the movement (Fig. 2 – Middle Left). Interestingly, many units in F5 were tuned for the hand used not only during the movement, but also from the end of the cue period onwards, showing a preference for either ipsi- or contra-lateral movements (Fig. 2 – Top Right). Additionally, units showing sustained tuning for grip were widely present in F5 (Fig. 2 – Middle Right), and occasionally units that were tuned for all three factors (Fig. 2 – Bottom Right).

**Figure 2.**
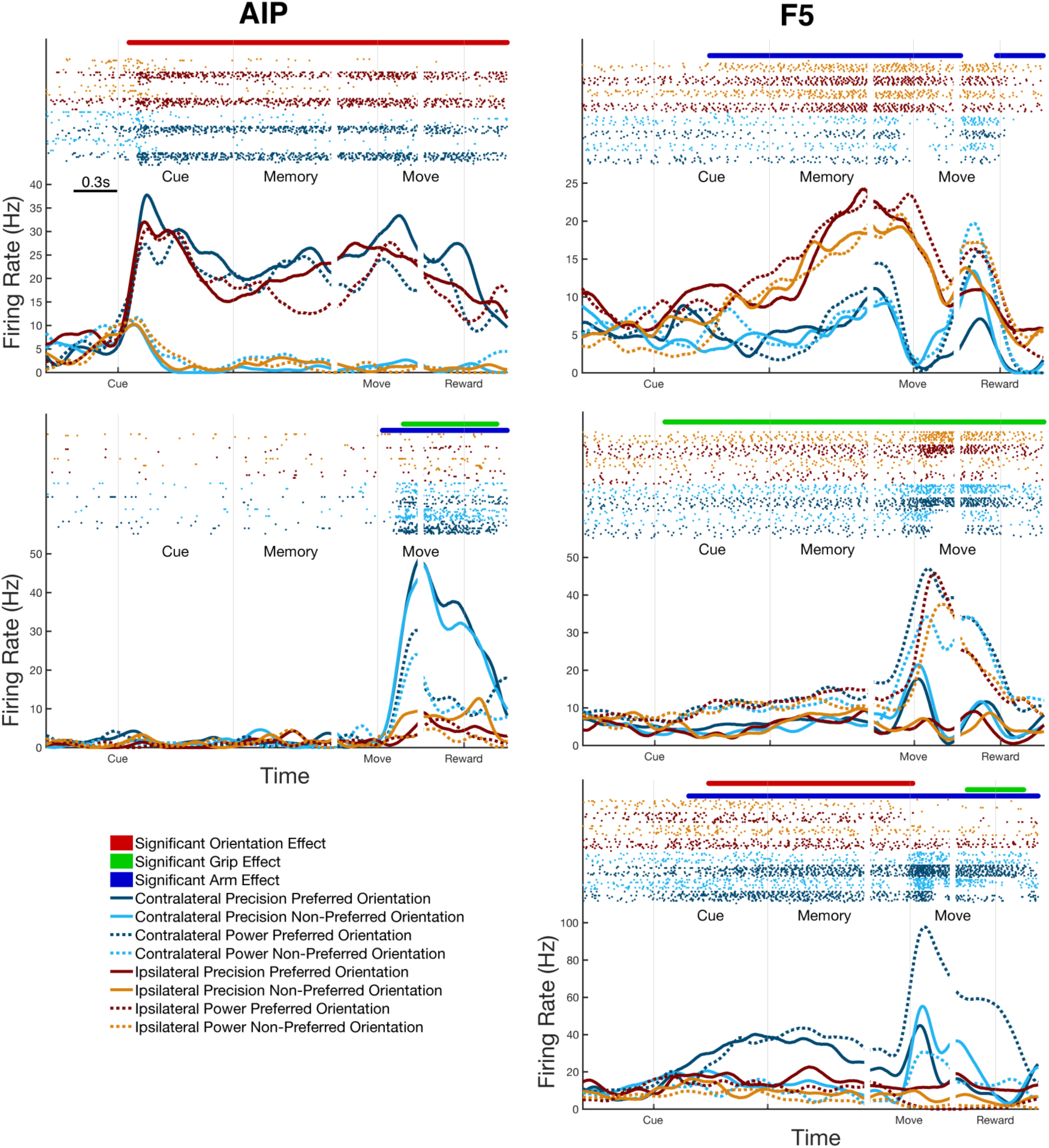
Example average firing rate curves of single-units in AIP and F5. (Top Left) A unit tuned to a single orientation of the handle throughout the trial, even in darkness. (Middle Left) A unit tuned only during movement both for the grip performed and the hand used. (Top Right) A unit tuned for hand used through the trial, showing a preference for ipsi-lateral movements. (Middle Right) A unit tuned for performed grip throughout the trial. (Bottom Right) A unit tuned for all task factors at different points in the trial. Data were aligned to three events, cue onset, movement onset, and reward. Raster plots above curves show single spikes over all trials of each condition. Significance bars represent tuning for each of the three factors, as determined by cluster-based permutation test (p < 0.05, corrected for number of factors). Examples were taken from both monkeys.

### Population response

Examining the average responses of individual units is an essential step. However, proper functional characterization requires examination of a population of units. While grip type and orientation tuning have been investigated in these areas previously, it is unclear how hand use affects these factors. Figure 3 shows the percentage of units tuned for grip or orientation, separately for contra- and ipsi-lateral trials. Orientation tuning was by far strongest in AIP and during the cue and movement periods for both monkeys. The amount of orientation tuning in F5 was low, especially for ipsi-lateral movements and in monkey P. Grip type tuning was highest during the movement period, and highest during the memory period highest for F5. In general, the amount of tuning for orientation and grip type was extremely similar for contra- and ipsi-lateral movement during movement planning, with the exception of orientation tuning in F5 in monkey S, where orientation tuning during ipsi-lateral trials decays quickly after the presentation of the cue. Although the amount of tuning differs between the monkeys in some cases, the results are qualitatively similar.

**Figure 3.**
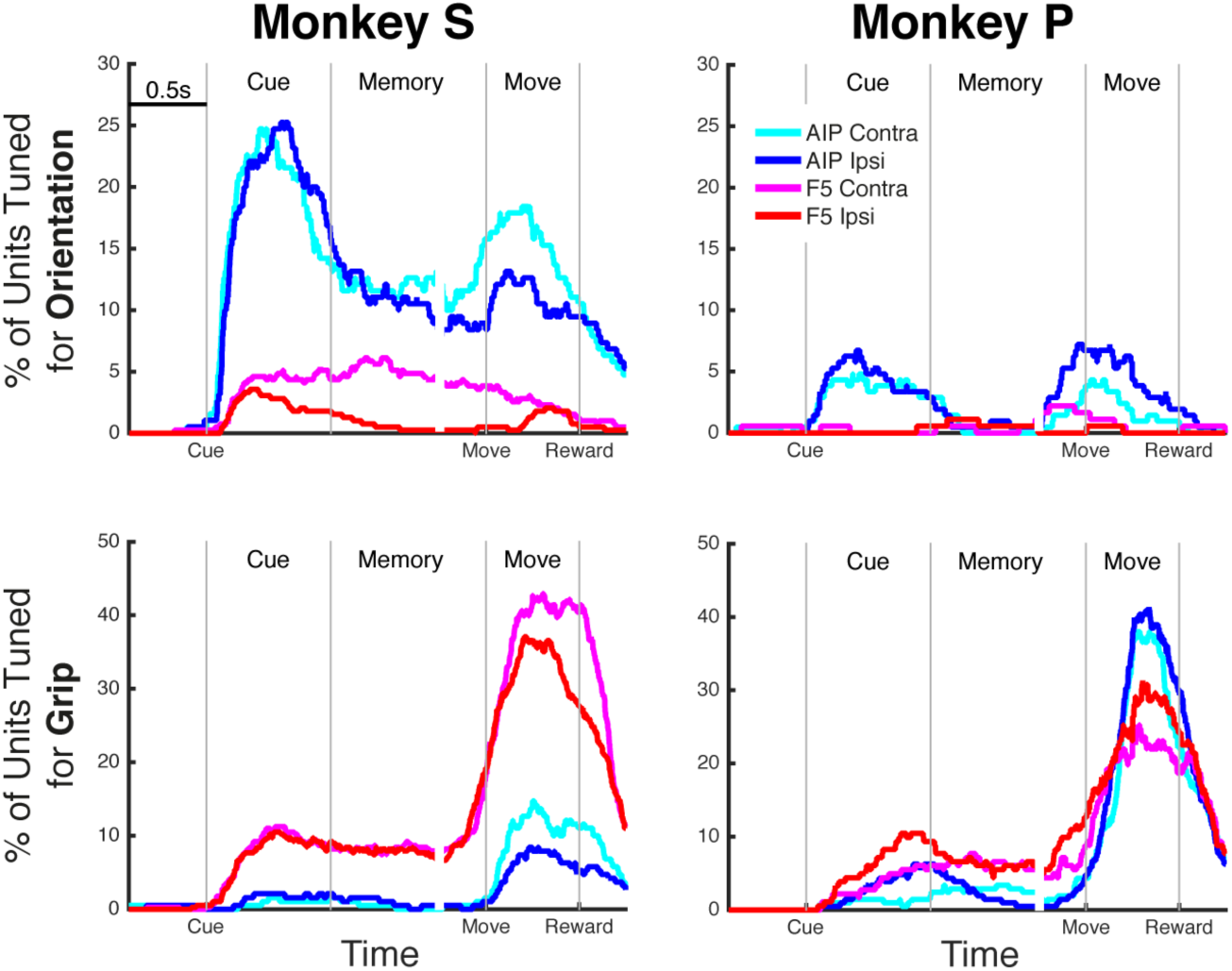
Grip type and orientation tuning over all recorded units, separated by hand used. Percentage of units tuned are plotted for orientation (Top) or grip type (Bottom) over time, separately for each monkey and as determined by the cluster-based permutation test (p < 0.05, corrected for the number of factors) run separately on trials of the contra- and ipsi-lateral hands. Data were aligned to three events: cue onset, movement onset, and reward.

Another interesting question is whether or not the preferred grip type was shared between contra- and ipsi-lateral grasps (i.e. was there an interaction between grip preference and hand preference). To test this, we compared the preferred grip type (highest firing) between trials of each hand for each unit that was significantly tuned (based on CBPT) during both movements. During cue and memory, not a single unit in either area switched grip type preference between contra- and ipsi-lateral trials, indicating that grip representation is completely shared regardless of hand used. However, during the movement period a portion of the units (∼5%) in both areas had differing grip preference between contra- and ipsi-lateral movements, a point that will be returned to in a later analysis.

### Laterality encoding

We have considered how hand use affects tuning for grip and orientation, but not the properties of hand tuning itself. Figure 4a shows the same CBPT analysis used previously for hand use. For monkey S, hand tuning was virtually non-existent in AIP before movement onset, suggesting that AIP encodes task-relevant features in a hand-independent manner before the movement has started. In contrast, hand tuning in F5 seemed to ramp continuously throughout the entire trial, reaching a maximum (50% of units tuned) just before the hold period, showing that F5 is strongly dependent on hand used. Tuning for hand use was high in both areas during movement itself, although more so in F5.

**Figure 4.**
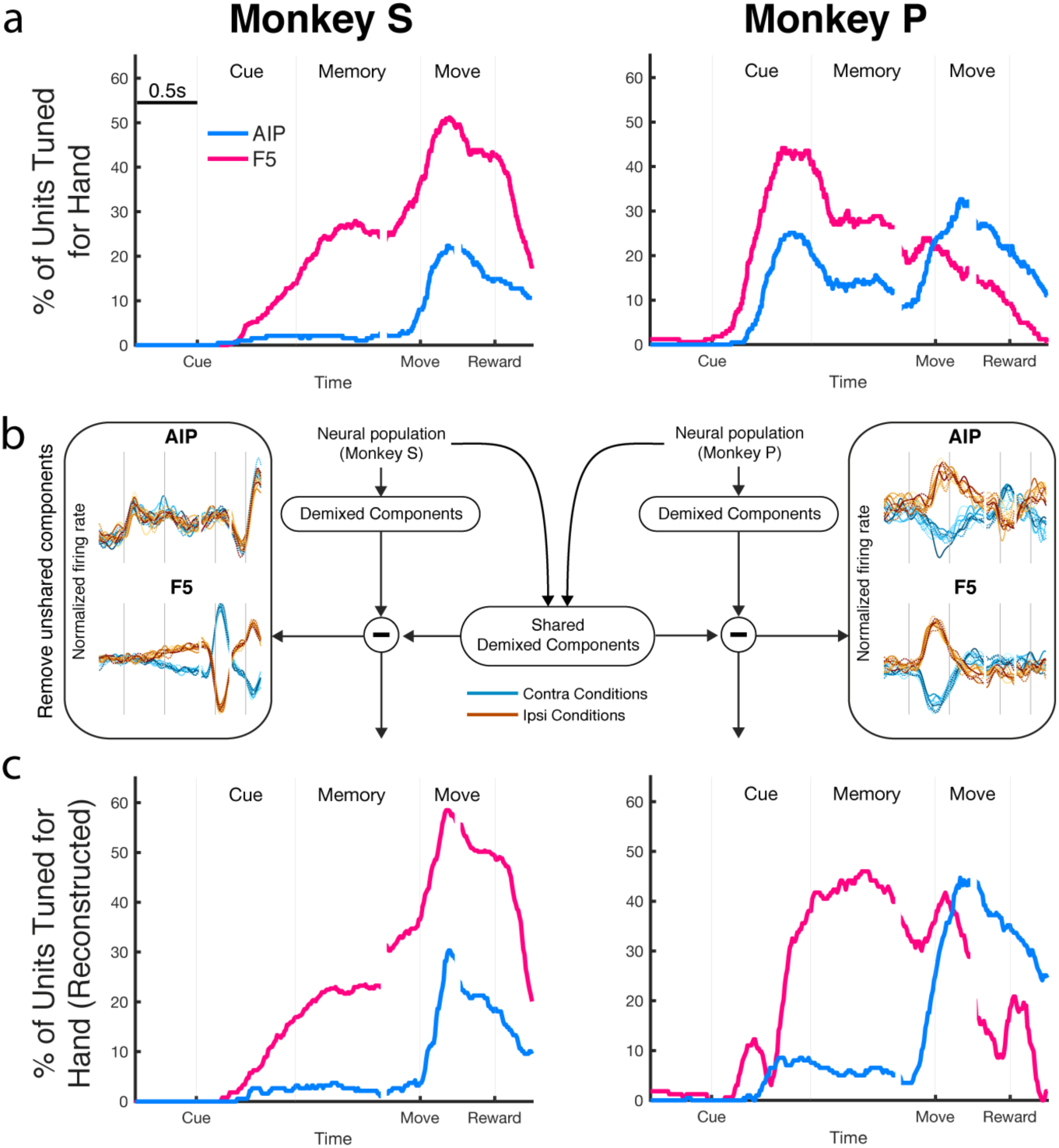
Extracting shared neural population components. (a) Percentage of units tuned for hand used, separately for each monkey and as determined by the cluster-based permutation test (p < 0.05, corrected for the number of factors). (b) Schematic of dimensionality reduction technique (dPCA) to demix neural population signals to find condition-specific projections that are shared between monkeys. dPCA is run on pooled data from both monkeys, separately per area, and dimensions that were not independently present in both monkeys are removed. Left-most and right-most panels show the most dominant removed dimensions (Methods). (c) After unshared dimensions are removed, the data of each monkey was projected back to the full neural space. Units were then tested again for hand tuning using a simple threshold matched to the baseline level of tuning in (a), revealing how the removal of unshared components affects the full neural space.

The results of monkey P with respect to hand tuning were significantly different, showing phasic bumps in hand tuning in both areas shortly after cue onset. As described in the Methods, a motion tracking program using an infrared camera tracked the position of the hands during the cue and memory periods to ensure that no premature movements occurred (strictly <1-2% change from baseline). However, given the differences between monkeys, we went back to the kinematic data to see if any biases in subthreshold movement could partially explain these differences. Using a linear classifier (leave-one-out cross-validated, Matlab function: *fitcdiscr*) on the recorded kinematics (Methods) during the memory period, it was possible to decode the intended hand on single trials with 52% accuracy in monkey S, where 50% is chance level, suggesting that no biases were present. In contrast, the hand used could be decoded with 75% accuracy in monkey P, specifically due to left hand trials, suggesting that there was a bias in subthreshold kinematics that could partially predict the intended hand. However, grip type could never be decoded from the infrared data during the memory period (50% accuracy in both monkeys). Taken together, these results suggest that monkey P may have made very small premature movements (1-2% change with respect to baseline) during the memory period that caused phasic spikes in hand tuning, but left other forms of tuning unaffected. Given the interesting finding that these differences in hand tuning didn’t seem to affect the other factors, we developed a method to extract factor-specific dimensions from the population of neurons that were shared between monkeys, described in the following section.

### Demixing shared population signals

As noted so far, both AIP and F5 are involved in the processing of a large multitude of task factors. These factors must be processed in parallel, and are distributed over many units in the population. To extract a population-level picture of these factors, we implemented a demixed principal component analysis (dPCA), a dimensionality reduction method for extracting low-dimensional linear combinations of neural populations that represent specific task features^29^. Since there was a significant difference in how hand was encoded between the two monkeys, but that didn’t affect the other task features, we developed an additional procedure for extracting only dimensions that were present independently in both monkeys (Methods), which is outlined in Figure 4b. The population data of each area was transformed into a set of dPCs, both for each monkey separately, and from a pooled set of neurons. The dPCs of each monkey were correlated with the shared components, and all components that had a correlation of at least 0.6 were retained, while the rest were discarded. The largest of these unshared components are shown on the side panels in Figure 4b. Interestingly, the largest components in both AIP and F5 of monkey P contained phasic tuning to hand during the cue, matching the period where monkey P likely made very small premature movements of the hand. By using the encoder found by dPCA (Eq. 2) to transform the retained dimensions back to the individual neural signals, and applying a threshold on the difference between the activity for trials of each hand, we can reconstruct the tuning of individual neurons (Fig. 4c). These results show that after this procedure the hand tuning in monkey P much more resembles the results for monkey S, showing a large amount of hand tuning in F5 during the memory period and almost none in AIP. It’s important to note that the reconstruction of the neural signal of each monkey rely only on data recorded from that animal, and therefore cannot artificially induce tuning, but only reveal structure that is present in the population signal.

Now that we’ve extracted the shared population signals, Figure 5 plots the shared dPCs over both monkeys. Thirty dimensions were extracted in each area, of which 17 and 20 were retained in AIP and F5, respectively. The excluded dimensions only accounted for 6% and 4% of the variance in the AIP and F5 data, respectively. As can be seen in Figure 5a,c, most of the variance of the neural population was captured in both areas (>80%), even after discarded non-shared dimensions. Intriguingly, the dPCA components taking up the most variance overall were condition independent signals (Fig. 5b). The largest component in AIP was a large condition-independent signal that modulates shortly before movement onset (component #1), while the second component was a large phasic response to the cue that also included a movement component. In F5, the largest component was a movement signal (Fig. 5d - component #1), while the second component was also mostly movement dominated. The next largest component in both areas was related to the hand used (component #3), although it was only possible to decode hand use from this signal in AIP during movement, as denoted by the black bars (Methods), while in F5 this was possible starting from early in the cue and maintained throughout the entire trial. Grip decoding was present in F5 throughout the trial starting in early cue (component #5), but most dominant in AIP during movement (component #4). Hand use and grip were also decodable from AIP before the movement (components #8 and #14), but accounted for a much smaller portion of the variance. Interestingly, handle orientation was very dominant in AIP throughout the trial, maintaining a consistent representation throughout the trial until after the reward, while less than one fifth as dominant in F5. The main results from this analysis suggest that population signals for task parameters can be demixed, and that hand and grip information are dominant in F5 during movement preparation, while orientation information is more dominant in AIP.

**Figure 5.**
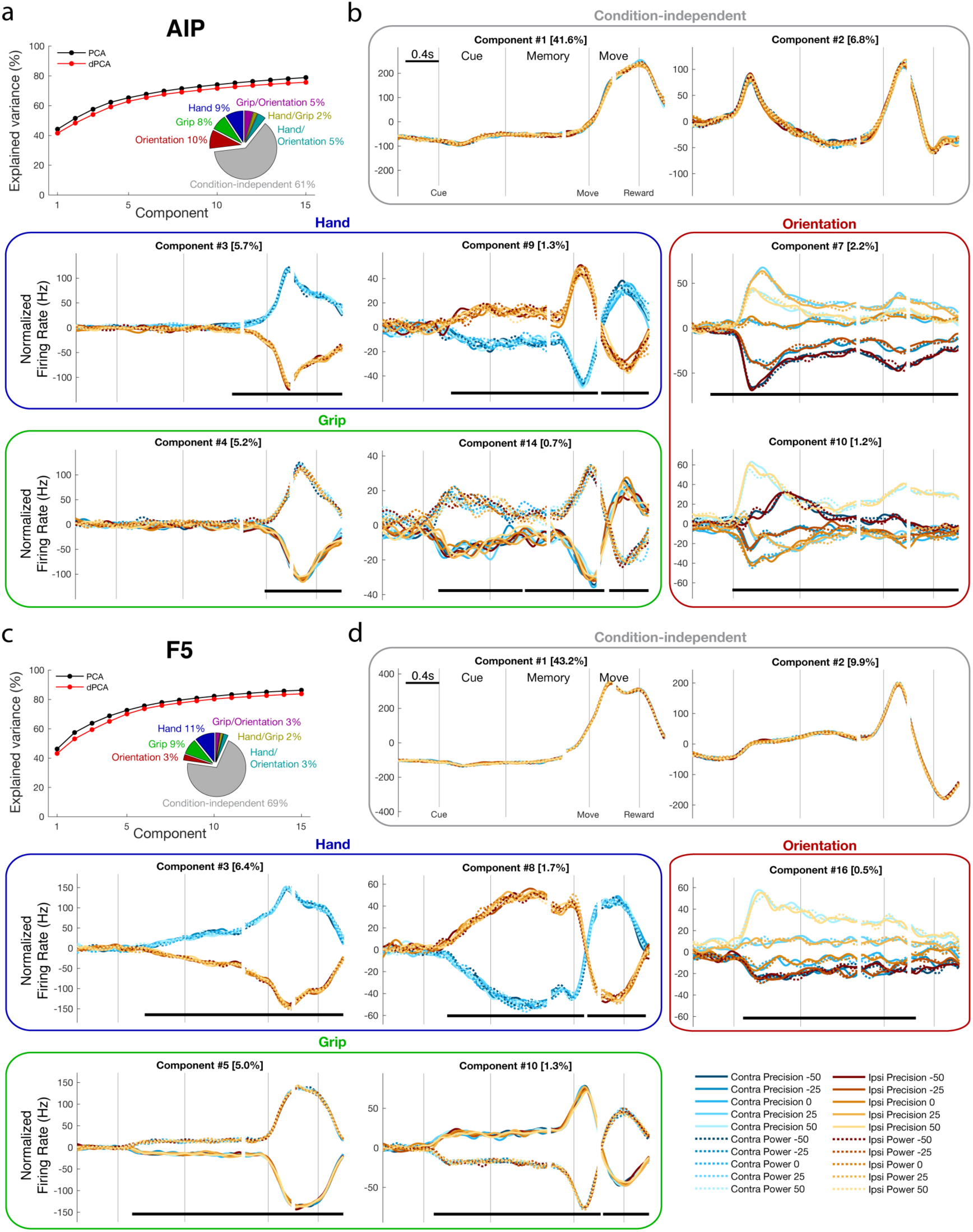
Demixing shared population signals. (a) Cumulative signal variance explained by PCA (black) and dPCA (red) in AIP. dPCA explains almost the same amount of variance as standard PCA. Pie chart shows how the total signal variance is split between dimensions. (b) Individual dPCs, separated into boxed by factor. Thick black lines show time intervals during which the respective task parameters can be reliably decoded from single-trial activity. Note that the vertical scale differs across subplots. (c-d) same as (a-b) for F5 data.

### Coordinate frame

Both areas investigated in the current study are essential parts of the visuo-motor transformation process, and therefore involved in transforming visual information into a body-centered coordinate frame so that muscle movements can be executed to the appropriate location in physical space. By examining how grip and orientation information changes between the hand used, it is possible to compare the representation of extrinsic (visual-centric) and intrinsic (body-centric) coordinate frames in the population.

Figure 6 illustrates dPCs that represent the interaction between the hand used and the other two task factors. Starting with grip type coding, no dimensions were found that showed a significant interaction of grip type and hand use before movement onset in either area (Fig. 6a), suggesting that grip representation is entirely independent of intended hand during movement planning. These results are in line with the earlier results that grip type preference was completely stable between hands during the cue and memory periods. Furthermore, these results suggest a visual-centric representation of grip type in both areas, since the visual cue is identical between trials of each hand. However, given that it is not clear how a body-centric coordinate frame should look for grip type, it is difficult to drawn firm conclusions.

**Figure 6.**
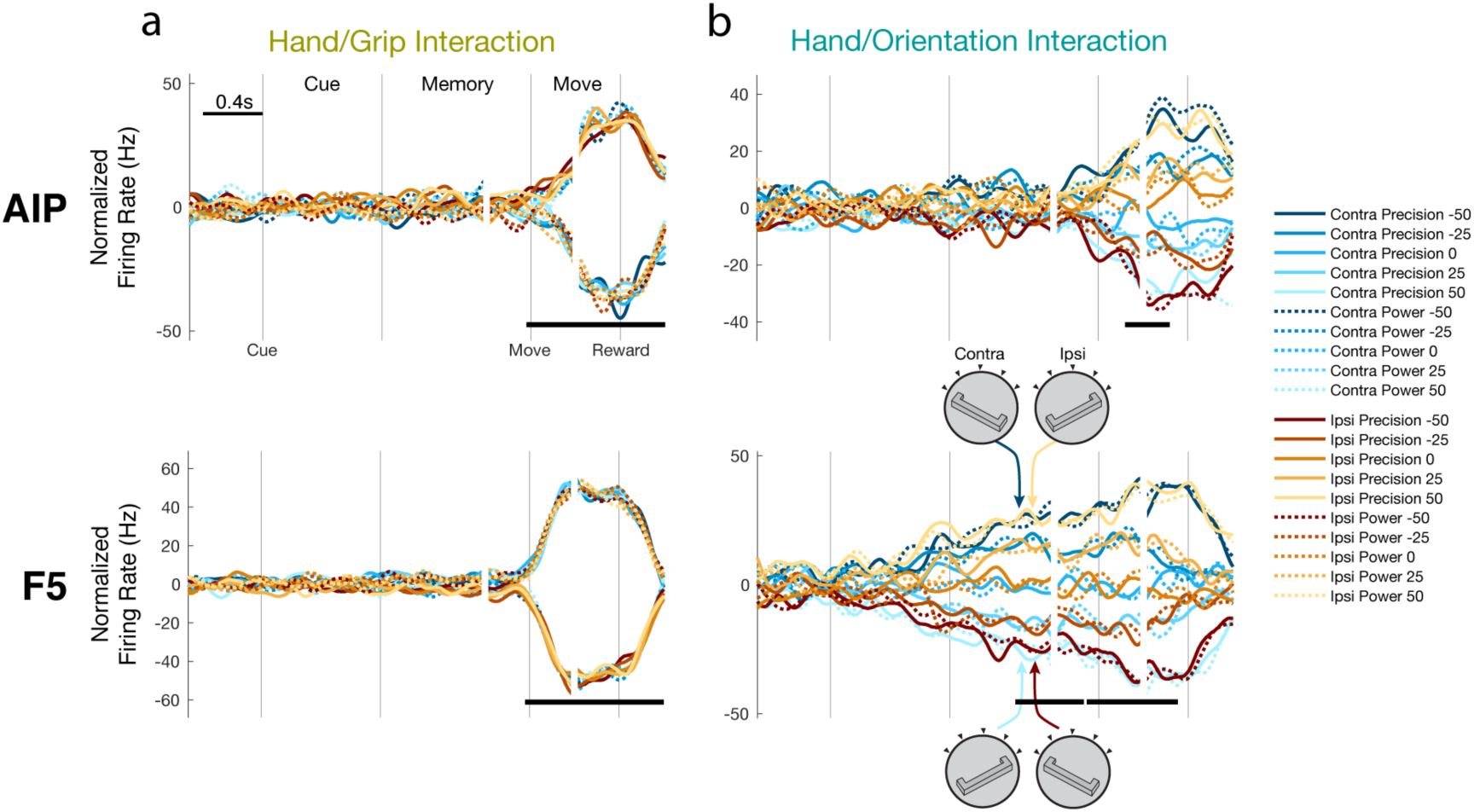
Interactions between hand use, grip type, and handle orientation. (a) One dimension was found with in both AIP and F5 showing a significant interaction between hand use and grip type, showing an interaction strictly after movement onset. (b) One dimension was found in both AIP and F5 showing a significant interaction between hand use and orientation. In both cases, the interaction showed a clustering of mirror-symmetric conditions (body-centric), as illustrated by the handle cartoons for the most extreme orientations. However, while this interaction was only present during late movement in AIP, while it was present starting from late cue in F5.

To further test visual-vs body-centric coordinate frames, we investigated the interaction between hand use and handle orientation. In a visual-centric frame, orientation representation should be shared regardless of hand used. In contrast, in a body-centric frame the orientation preference may shift between hands to match the correspondingly mirrored wrist rotation (i.e., amount of pronation or supination) required between each hand. For example, analogous muscle contractions are produced in the contra- and ipsi-lateral hands during grasping of a handle oriented in a mirror-symmetric fashion (e.g. - 50° matched with a 50° rotation). Interestingly, we found that during late movement mirror-symmetric conditions clustered together in AIP (Fig. 6b), which could represent sensory feedback relayed to parietal cortex. Crucially, we found a similar dimension in F5 that began showing mirror-symmetric clustering starting shortly after the cue and maintaining the same representation throughout the movement, showing a clear presence of a body-centric coordinate frame during movement planning. It’s important to note that nothing in our analysis method enforces that mirror-symmetric clustering appear in the interaction component as opposed to any other kind of interaction between contra- and ipsi-lateral movements. Taken together, these results show that while grip representation is independent of hand use before movement onset, there exists a body-centric coordinate frame for wrist orientation in F5 during movement planning.

## Discussion

In the current study, by recording from populations of neurons in AIP and F5 of two macaque monkeys during a delayed grasping task, we found that the laterality of hand use was only weakly represented in AIP before movement, while intended hand use was dominant in F5. Furthermore, grip representation was independent of hand use before movement, as was much of the orientation information, which was far more dominant in AIP. However, a small portion of the neural population in F5 showed representation of handle orientation in a body-centric coordinate frame during movement preparation and into movement, implicating a visuo-motor process taking place between AIP and F5.

It’s interesting that in the current study grip preference and tuning was identical regardless of hand use. It has been found that after severing all connections between hemispheres of macaques the ipsi-lateral hemisphere can coordinate reaches, but not properly pre-shape the fingers of the hand to grasp food^15^. Based on our results, premotor cortex in either hemisphere should have the required information for successful hand shaping. Therefore, it is unlikely that involvement of F5 in ipsi-lateral movements is contributing directly to muscle activation. Supporting this, stimulation of M1 produces no direct corticospinal activation of ipsi-lateral muscles^30^. Additionally, a rodent study has shown that although bi-lateral representation is common in premotor cortex, neurons projecting to the brainstem showed mostly contra-lateral preference^31^. Therefore, F5 modulation during ipsi-lateral movements is likely coordinated with the contra-lateral F5 through the corpus callosum and likely plays a larger role during bi-manual rather than uni-manual grasping movements, as is the case in M1^32^.

Interestingly, while grip preference did not change between hands, many units changed their grip or hand preference between the preparatory periods and the movement itself (Fig. 5), in line with studies showing that individual unit tuning tends to be unstable, and importantly that different dynamics govern preparation and movement^33-35^. The amount of grip tuning obtained in AIP was significantly lower than found in previous studies^20^, as was the amount of orientation tuning found in F5^21^. We believe these differences are due to selective recording of task-related units in previous studies, while in the current study units were not evaluated for tuning online, presumably giving a more unbiased estimate of tuning percentage.

The fact that AIP showed very little preparatory response to hand use is unexpected, especially since the nearby parietal regions involved in reaching show strong modulation^1-3^. Additionally, AIP is part of the network that responds to passive auditory listening^36^, and since the current task employed an auditory cue, it would be expected that playing differing tones cuing hand use would elicit a task dependent response. It could be that the auditory stimuli in our task were not varied enough to elicit a significant effect, or rather that since the task was active rather than passive, AIP was likely dominated by visual processing demands. During the movement itself both areas strongly represented the hand used. Along with the established role of F5 in ongoing movement generation, hand tuning during movement could originate from projections from secondary sensory cortex to both F5^37,38^ and AIP^39^. Although monkeys received grip cue information at the center of their visual field, the effector cue was auditory, introducing a potential confound in lateralized processing. However, it is unlikely that any lateralization effects found in the current study are a result of asymmetric processing of auditory information since only complex stimuli, such as vocalizations, evoke a lateralized response in macaque monkeys^40,41^.

Since AIP showed almost no hand-specificity before movement, it is unlikely that the preparatory hand tuning observed in F5 originates from AIP. The hand tuning in F5 likely comes indirectly through prefrontal cortex, from which a number of areas project to F5^37^, also in line with the fact that hand tuning in F5 appears only towards the end of the cue, as observed previously^6^. An alternative explanation for large amounts of hand-invariant tuning could be that many of the same proximal muscles are required for movements of either arm, given the large postural adjustments required in extending the arm. However, experiments limiting movements to the distal muscles alone^10^, or controlling for postural contributions to ipsi-lateral control^42^, have shown a strongly bi-lateral representation of hand movement in premotor cortex, suggesting that postural control cannot fully explain hand-independence.

Based on our analysis of infrared motion tracking, it is very likely that monkey P made small movements of the hand during the preparatory phases, biasing hand tuning during that time. However, the same analysis showed that no such movements occurred with monkey S, and grip and orientation tuning appeared unaffected in both monkeys. Furthermore, we were able to extract the population level preparatory and movement related signals that were shared in both monkeys (Fig. 5), revealing the commonality in data sets. A number of studies support the notion that low-dimensional features of neural populations have a biological basis, including learning in brain-computer interfaces^43^, gating of motor output^44^, and parallel encoding^45,46^.

Monkeys were required to reach to the target as well as grasp. Therefore, reach planning and execution is likely a significant part of the observed activity. However, as we have argued previously^25^, previous research employing a grasp-only task^47^ and a grasp-reach dissociation task^48^ suggests that F5 encodes grasping quite independently of reaching, although both areas contain information about reach position^49^. Furthermore, reversibly inactivating F5^50^ or AIP^51^ selectively impairs hand-shaping and not reaching, suggesting that our results are an accurate representation of the grasping network.

While visual-centric coordinate frames were present in AIP and F5, a body-centric frame was only found in F5. Finding both representations in F5 is not altogether surprising, since the ventral premotor cortex, as well as dorsal premotor cortex^9^, has been shown to be very sensitive to visuo-spatial information as opposed to the dynamics of movement^6,52^. However, ventral premotor cortex likely shifts its control strategy during movement^53^. F5 is therefore likely a site of transformation between hand-invariant and hand-specific representations, representing stimuli from both the contra- and ipsi-lateral visual hemi-field^54,55^, leading to a visuo-spatial dependence. Overall, the presence of a body-centric coordinate frame in F5 during preparation reveals a more direct representation of sensorimotor integration than posited previously^56^, and provides a novel perspective on the functional properties of the parieto-frontal grasping circuit.

## Methods

### Experimental Setup

Two female rhesus monkeys (Macaca mulatta) participated in this study (monkeys P and S; weight 4.5 and 5.5 kg, respectively). They were pair-housed in a spacious and enriched environment. All procedures and monkey care were conducted in accordance with the regulations set by the Guidelines for the Care and Use of Mammals in Neuroscience and Behavioral Research^57^, and in agreement with German and European laws governing monkey care.

Monkeys were habituated to comfortably sit upright in an individually adjusted primate chair with the head rigidly fixed to the chair. A grasp target was located at a distance of 24 cm in front of the monkey. The target consisted of a handle that could be grasped with two different grip types, either with a precision grip (using index finger and thumb in opposition) or a whole-hand power grip^20,21^. Inside the handle, two touch sensors were placed in small, visible recessions to detect the contact of the monkey’s thumb and index finger during precision grips. An infrared light barrier placed inside the opening of the handle detected power grips. Grip type was instructed by two colored light emitting diodes (LEDs) that were positioned immediately above the grasping handle. The handle was rotatable and was presented in five different orientations (upright and 25° or 50° clockwise and counter-clockwise) and two spotlights could illuminate it from the left and right side in an otherwise dark experimental room. Two capacitive touch sensors (model EC3016NPAPL; Carlo Gavazzi) were placed at the level of the monkey’s waist as handrest buttons. A single speaker, which produced the audio tones for cuing the appropriate arm, was positioned directly above and behind the monkey’s head. The speaker was oriented such that the audio tone was equally directed into each ear. Monkeys had to fixate on a red LED that was positioned between the two cue LEDs. Eye movements were measured using an optical eye tracker (ET-49B; Thomas Recording) and custom-written software implemented in LabView Realtime (National Instruments) using a time resolution of 5 ms was used to control the behavioral task.

### Behavioral paradigm

Monkeys were trained in a delayed grasping task in which they were required to grasp a handle in five possible orientations with either a power grip or a precision grip using the left or right hand. This led to 20 grasp conditions that were presented in a pseudorandom order. To initiate a trial, monkeys sat in darkness and placed each hand on a handrest button. The handle was then positioned in one of the five orientations over a fixed duration (approximately 2 seconds) and subsequently a red fixation LED switched on. From then on, the monkey was required to fixate while keeping both hands still (see Hand motion tracking section) on the handrest buttons (fixation period duration: 700–1100 ms, mean: 900 ms), as illustrated in Figure 1a,b. In the following cue period (cue period duration: 800 ms), the object was illuminated to reveal its orientation. The color of an additional LED presented to the left or right of the fixation LED indicated which grip type to perform, either a power grip (green light, left) or a precision grip (yellow light, right). In addition, an audio tone (1000Hz or 2000Hz), representing the left and right arms, respectively, was presented simultaneously with the grip cue and spotlights. The spotlights, audio tone, and the grip cue LED were then switched off while the fixation light remained on for a variable period (memory period duration: 700–1100 ms, mean: 900 ms) during which the monkey had to remember the trial instructions. A brief blinking of the fixation LED (130 ms) instructed the monkey to reach and grasp the object in the dark with the correct arm while maintaining eye fixation and keeping the other arm on the handrest. After a hold period of 300 ms, each correct trial was rewarded with a fixed amount of water.

### Hand motion tracking

In addition to normal behavioral control, the stationarity of each monkey’s hands on the hand rests was also continuously tracked during the late cue and memory period of every trial with an infrared camera positioned directly over the hands in combination with a separate custom-written LabView control program. In detail, at every time point a reference image that was recorded before training (view of the handrest without monkey present) was subtracted from the current view of the hands and this image was subsequently thresholded in order to isolate the infrared data specific to the hands themselves. In this way, the stationarity of both hands could be simultaneously monitored for several criteria: (a) the total luminance of the hand, (b) the center of the hand, i.e. the position of the weighted average of the most luminous pixels in both the x and y direction separately, and (c) the standard deviation around the center of the hand in both the x and y direction. During the first 400 ms of the cue period on every trial the value of each of these measures was recorded as a reference. If, at any subsequent time before the go cue, either hand deviated ±1-2% with respect to the reference values recorded during the early cue, the trial was aborted without reward. The thresholds beyond which a trial would be aborted were fixed for each of these three factors during all recordings of both monkeys at: (a) ±2%, (b) ±1%, and (c) ±2%.

Additionally, during all recordings of monkey S and a portion of the recordings from monkey P, continuous infrared hand motion information was digitally stored (500 Hz) for additional offline analysis. For each hand the sum of a portion of the above-mentioned variables was recorded, i.e. the average of the percent deviation in total hand luminance, the center of the hand in x-coordinates, and the center of the hand in y-coordinates from the reference values recorded during the beginning of the cue period of each trial. The storage of this data allowed for an offline analysis of sub-threshold movements.

### Surgical procedures and MRI scans

Details of the surgical procedures and MRI scans have been described previously^48^. In short, a titanium head post was secured in a dental acrylic head cap and a custom made oval-shaped recording chamber (material PEEK [polyether ether ketone]; outer dimensions, 40 X125 mm^2^; inner dimensions, 35 X 20 mm^2^) was implanted over the right or left hemisphere to provide access to parietal area AIP and premotor area F5.

Two structural magnetic resonance image (MRI) scans of the brain and skull were obtained from each monkey, one before the surgical procedures, to help guide the chamber placement, and one after chamber implantation to register the coordinates of the chamber with the cortical structures (Fig. 1c,d). AIP was defined as the rostral part of the lateral bank of parietal sulcus^39^, whereas recordings in F5 were made primarily in F5a and F5p, which are in the post-arcuate bank lateral to the tip of the principal sulcus^58^.

### Neuronal recordings

Single-unit and multi-unit (spiking) activity was recorded using quartz-glass-coated platinum/tungsten single electrodes (impedance 1–2 MΩ at 1 kHz) or tetrodes (impedance 500-800 kΩ at 1 kHz) that were positioned simultaneously in AIP and F5 by two five-channel micromanipulators (Mini-Matrix, Thomas Recording). Neural signals were amplified (400X), digitized with 16-bit resolution at 30kS/s using a Cerebus Neural Signal processor (Blackrock Microsystems), and stored on a hard drive together with the behavioral data.

### Preprocessing

All data analysis was performed offline. Neural signals were band-pass filtered (forward-backward) with cutoff frequencies between 300-5000 Hz. Waveforms were extracted when the signal deflected beyond 5 standard deviations from baseline either negatively or positively. The refractory period between spikes was set at 1.5 ms. During tetrode recordings spikes that were detected on one of the electrode tips were extracted from all 4 and aligned to the peak or valley of the first channel to cross the threshold. Units were isolated using principal component analysis techniques (Offline Sorter v3.2.2, Plexon), and sorted into single- and multi-units based on the inter-spikes interval, magnitude of spike waveform, and consistency of waveform shape. Using Matlab (Mathworks) for further analysis, we included all units in our database that were stably recorded for at least 5 trials per condition (100–260 trials in total). Average firing rate curves were generated using a Gaussian window as a kernel (SD: 57 ms) in three alignments (cue, movement, and reward).

### Data analysis

The preferred and non-preferred orientations were determined for each unit from the mean activity in the time interval from cue onset to reward onset. Activity was averaged across all trials of the same orientation. Of the five tested orientations, the orientation with the higher (or lower) mean firing rate was defined as the preferred (or non-preferred) orientation, as in Baumann et al.^20^.

To complement each firing rate curve, periods of significant tuning were calculated using a cluster-based permutation test (CBPT) to generate the significance bars in Figure 2^59^. Briefly, this test evaluates the t-statistic (independent samples) between two conditions over all time points and extracts clusters (consecutive time segments) of activity whose t-statistic exceeds a predefined threshold (*α* = 0.05), then the absolute t-statistics within each cluster were summed to produce cluster-level statistics. To generate a chance-level distribution from which the appropriate threshold could be determined, trials were randomly partitioned between the two conditions and the t-test and clustering redone (1000 partitions). From every partition the largest cluster-level statistic was used to generate a largest chance cluster distribution. By comparing the real cluster-level statistic against the largest chance cluster distribution, significant clusters could be determined if the observed cluster value exceeded a set percentage of largest chance cluster values (p = 0.05). In this way, sensitivity to short time-scale differences is greatly reduced, but the overall false-alarm rate remains below the designated p-value. This test was carried out once for each of the three factors. Additionally, to see if grip and orientation tuning differed between contra- and ipsi-lateral trials, the CBPT was repeated for those trials separately.

A unit was considered task-related if it had a significant effect of any of the three factors at any tested time point of the CBPT. Crucially, all analyses only considered units that were determined to be task-related. As a control, if a 3-way ANOVA is used in place of the CBPT, approximately the same amount of significance is found overall, suggesting that the CBPT does not over-estimate the level of tuning for each unit.

### Dimensionality reduction

A common problem with large data sets is their inherent complexity. Principal component analysis (PCA) is commonly employed to reduce the dimensionality of such data sets by finding a low dimensional representation of the data by defining independent linear combinations, or weighted averages, of units that explain most of the variance in firing rates. PCA finds a ‘decoder’, **D**, which represents a linear mapping of the full data onto a compressed read out. Using an ‘encoder’, **F**, data can then be approximately reconstructed by decompressing it.

To formalize this, given a matrix of data **X**, where each row contains the average firing rates of one neuron for all task conditions, PCA finds an encoder, **F**, and an equivalent decoder, **D**, which minimizes the loss function

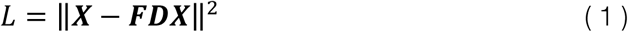

under the constraint that the principal axes are normalized and orthogonal, and therefore **D** = **F^T^**, with the Frobenius norm 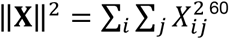^60^. Unfortunately, data that is represented in this way heavily mixes the effect of different task parameters between latent dimensions, since no information regarding the actual task conditions is present in the loss function.

However, we would like to extract dimensions that dissociate our specific task conditions. To achieve this, demixed principal component analysis (dPCA) was performed^29^ using freely available code: http://github.com/machenslab/dPCA.

dPCA is similar to classical PCA in the sense that it seeks to find a rotation of the full neural space that explains most of the variance in average firing rates in a small number of latent dimensions. In contrast to PCA, dPCA seeks to explain marginalized variance with respect to our specific task variables (hand, grip type, orientation, interactions, and time), instead of merely explaining total variance. The differences between traditional PCA and dPCA can be formalized by comparing the loss functions that are minimized in each procedure. dPCA utilizes a separate encoder and decoder

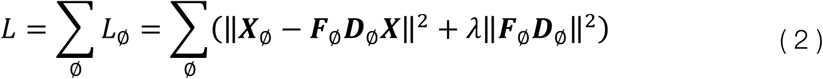

where **X**_*Ø*_ is the marginalization of the full data with respect to each of our task parameters of interest and the λ term is a regularization parameter, preventing overfitting. Marginalizations of **X** can be obtained by averaging over all parameters which are not being investigated and subtracting all simpler marginalizations (then replicating matrix entries so that **X**_*Ø*_ has the same dimensionality as **X**). In our case the marginalizations of interest were Time, Hand x Time, Grip x Time, Orientation x Time, Hand x Grip x Time, Hand x Orientation x Time, and Grip x Orientation x Time^61^. The value of λ was 5.1·10^-7^ and 2.3·10^-7^, respectively, as determined via cross-validation for the brain areas AIP and F5 in the data combined over both monkeys. There was only one value of λ used per brain area. All extracted dimensions were permitted to vary along the entire time axis in addition to their respective dimension.

In addition to finding demixed latent dimensions, our goal was to find latent dimensions in the pooled data of both monkeys that accurately represented aspects of the task that were present of both monkeys. Crucially, we wanted to exclude dimensions that could only explain variance in the units taken from a single monkey. In order to achieve this, dPCA was first carried out (with cross-validated regularization parameters) on the data of each monkey separately. Next, for each brain area, all dPCs of each individual monkey were correlated to the pooled dimensions. If any pair of dimensions produced an correlation of at least 0.6, those dimensions were considered to be robust in both monkeys, and all other dimensions were discarded.

A decoding procedure was undertaken to determine if the dPCs provided useful decoding axes for the task conditions. We used 100 iterations of stratified Monte Carlo leave-group-out cross-validation, where on each iteration we held out one random trial for each unit in each condition forming *X*_*test*_ (as the units were not recorded simultaneously, we do not have recordings of all units in any actual trial). Remaining trials were averaged to form a training set *X*_*train*_. We then calculated dPCA on *X*_*train*_ and used the resulting components as linear classifiers for the trials in *X*_*test*_. We then used 100 shuffles to compute Monte Carlo distribution of classification accuracies expected by chance. For each unit and iteration, we shuffled all available trials between conditions, respecting the number of trials per condition. If the real classification accuracy exceed that expected by chance on all iterations and for 200 ms contiguously, classification was considered significant and marked as black bars in Figures 5 and 6^29^.

## Acknowledgements

We would like to thank N. Bobb, R. Lbik, and M. Dörge for technical assistance, B. Dann for constructive discussions, and B. Lamplmair and S. Schaffelhofer for providing illustrations.

## Author Contributions

JAM and HS designed research; JAM performed research; JAM analyzed data; JAM wrote the paper; JAM and HS edited the manuscript.

